# Congruent population genetic structures and divergence histories in anther-smut fungi and their host plants *Silene italica* and the *S. nutans* species complex

**DOI:** 10.1101/2020.01.04.894774

**Authors:** Fanny E. Hartmann, Alodie Snirc, Amandine Cornille, Cécile Godé, Pascal Touzet, Fabienne Van Rossum, Elisabeth Fournier, Stéphanie Le Prieur, Jacqui Shykoff, Tatiana Giraud

**Author notes:** Author for Correspondence: Fanny E. Hartmann, Ecologie Systematique Evolution, Bâtiment 360, Univ. Paris-Sud, AgroParisTech, CNRS, Université Paris-Saclay, 91400 Orsay, France.

## Abstract

The study of population genetic structure congruence between hosts and pathogens gives important insights into their shared phylogeographic and coevolutionary histories. We studied the population genetic structure of castrating anther-smut fungi (*Microbotryum* genus) and of their host plants, the *Silene nutans* species complex, and the morphologically and genetically close *S. italica,* which can be found in sympatry. Phylogeographic population genetic structure related to persistence in separate glacial refugia has been recently revealed in the *S. nutans* plant species complex across Western Europe, identifying several distinct lineages. We genotyped 171 associated plant-pathogen pairs of anther-smut fungi and their host plant individuals using microsatellite markers and plant chloroplastic SNPs. We found clear differentiation between fungal populations parasitizing *S. nutans* and *S. italica* plants. The population genetic structure of fungal strains parasitizing the *S. nutans* plant species complex mirrored the host plant genetic structure, suggesting that the pathogen was isolated in glacial refugia together with its host and/or that it has specialized on the plant genetic lineages. Using random forest approximate Bayesian computation (ABC-RF), we found that the divergence history of the fungal lineages on *S. nutans* was congruent with the one previously inferred for the host plant and likely occurred with ancient but no recent gene flow. Genome sequences confirmed the genetic structure and the absence of recent gene flow between fungal genetic lineages. Our analyses of host-pathogen individual pairs contribute to a better understanding of co-evolutionary histories between hosts and pathogens in natural ecosystems, in which such studies are still scarce.

## Introduction

Host-pathogen interactions are pervasive in natural ecosystems, with many important ecological and evolutionary consequences (Poulin, 2005; Thompson, 2005). Pairs of tightly interacting hosts and pathogens may share common evolutionary histories due to co-evolution and/or shared geographic and climatic constraints, or may follow distinct evolutionary trajectories due to differences in reproductive systems, dispersal ranges, population sizes or contingent histories (Tellier, de Vienne, Giraud, Hood, & Refrégier, 2010; Thompson, 2005). The comparison of genetic divergence histories between host and pathogen populations gives important insights into their shared phylogeographic history and possible local adaptation or host specialization (Croll & Laine, 2016; Feurtey et al., 2016). Studies of phylogeny congruence of hosts and pathogens can for example allow identifying host shifts or co-speciation events (de Vienne et al., 2013; Hafner & Page, 1995; Wilson, Falush, & McVean, 2005). Several cases of host shift or co-speciation events between hosts and pathogens have led to damaging diseases in plants, animals and humans (Fisher, Gow, & Gurr, 2016; McDonald & Stukenbrock, 2016; Wolfe, Dunavan, & Diamond, 2007). The study of population genetic structure congruence within species or species complexes can help identifying and understanding patterns of local adaptation (Gandon, Capowiez, Dubois, Michalakis, & Olivieri, 1996), which play an important role in the dynamics of pathogen and host communities (Gandon & Michalakis, 2002; Laine, 2005; Laine, 2008). Studies investigating the congruence of population genetic structure and divergence histories between hosts and pathogens are yet surprisingly still scarce despite their importance for understanding the evolutionary mechanisms and histories leading to host specialization and local adaptation (Barrett, Thrall, Burdon, & Linde, 2008; Croll & Laine, 2016), but see (Dybdahl & Lively, 1996; Feurtey et al., 2016; McCoy, Boulinier, & Tirard, 2005; Michalakis, Sheppard, Noel, & Olivieri, 1993; Tsai & Manos, 2010).

Anther-smut fungi (*Microbotryum* genus) are generally highly specialized on their host plant species of the Caryophyllaceae family (Hartmann et al., 2019; Kemler, Göker, Oberwinkler, & Begerow, 2006; Le Gac, Hood, Fournier, & Giraud, 2007; Refrégier et al., 2008) and patterns of local adaptation have been reported in the system (Feurtey et al., 2016; Kaltz, Gandon, Michalakis, & Shykoff, 1999). Therefore, they constitute a highly suitable system to study the congruence of population genetic structure and divergence history between hosts and pathogens at different evolutionary scales. Comparisons of phylogenies at the genus level suggested a prevalence of host shifts at large evolutionary scales (Refrégier et al., 2008). The plant *Silene latifolia* and its anther-smut fungus *M. lychnidis-dioicae* display strong congruence of population genetic structures and plant local adaptation at regional and continental scales, suggesting the existence of co-evolution in the system (Delmotte, Bucheli, & Shykoff, 1999; Feurtey et al., 2016; Kaltz et al., 1999). Population genetic structures of both the host and the fungal pathogen likely resulted from past climatic events, showing hallmarks of recolonization from former glacial refugia in Europe (Badouin et al., 2017; Gladieux, Devier, Aguileta, Cruaud, & Giraud, 2013; Gladieux et al., 2011; Taylor & Keller, 2007; Vercken et al., 2010). The congruence of host and pathogen population genetic structures has not been investigated in other anther-smut fungi-*Silene* pairs despite their importance as models of pathosystems in natural ecosystems (Bernasconi et al., 2009; Toh & Perlin, 2016) and the importance of assessing whether congruence in population genetic subdivision is a general pattern (Croll & Laine, 2016). Furthermore, the evolutionary history of population genetic divergence and of gene flow levels occurring between pathogens infecting closely related hosts with overlapping distribution ranges still remain poorly studied at the genome-wide-scale (but see Badouin et al., 2017).

*Silene nutans* is an assemblage of strongly differentiated cryptic genetic plant lineages, corresponding to at least seven “evolutionary significant units”, which can be grouped into two main phylogeographic (eastern and western) genetic clusters in Europe (Van Rossum et al., 2018). The *S. nutans* genetic lineages display several morphological and ecological differences, as well as nuclear and plastid genetic differentiation (Martin et al., 2016; Van Rossum et al., 2018), and strong postzygotic reproductive isolation has been shown between the eastern and western clusters in Western Europe (Martin et al., 2017). Geographic distribution patterns suggest that *S. nutans* lineages have diverged in allopatry during the Quaternary climate oscillations, and then recolonized northwards, without admixture (Martin et al., 2016; Martin et al., 2017; Van Rossum et al., 2018). The anther-smut fungal species *Microbotryum violaceum sensu stricto* is specialized on *S. nutans* (Kemler et al., 2006; Lutz et al., 2005). *Microbotryum* fungi are pollinator-borne pathogens castrating plants by replacing the pollen by their own spores and aborting ovaries, and they are usually highly specific on their *Silene* host plant species (Le Gac, Hood, & Giraud, 2007; Refrégier et al., 2008). The existence of cryptic lineages within *S. nutans* therefore raises the question of whether cryptic lineages also exist in the fungus and whether their genetic divergence history mirrors that of the host. In Southern Europe, *S. nutans* can be found in sympatry with *S italica*, a closely related species, in particular in the Cévennes (Lafuma & Maurice, 2006). *Silene nutans* and *S. italica* can be difficult to distinguish, being very similar morphologically, mainly differing in petal shape, in the length of the gynophore and with rather nodding or erect flowers, respectively (Rameau, Mansion, & Dumé, 1989, 2008; Tison & de Foucault, 2014). Their flowering time, pollinator guilds and ecological niches overlap, although they slightly differ in their ecological requirements, with *S. italica* being strictly calcicolous, more xero-thermophilous and heliophilous than *S. nutans* (Rameau et al., 1989, 2008; Tutin et al., 2001). *Microbotryum* fungi have been found on the two plant species and appeared differentiated based on a small sample and a few genetic markers (Bucheli, Gautschi, & Shykoff, 2000); however, spillover, i.e., non-sustainable cross-species disease transmission, has been shown to occur in *Microbotryum* fungi (Antonovics, Hood, & Partain, 2002; Gladieux et al., 2011). The study of the population genetic structure of *M. violaceum s. s.* on *S. nutans* therefore requires molecular typing to check plant and fungal species identities.

In this study, we analyzed associated plant-pathogen samples of *S. nutans* and *S. italica* populations from Europe in order to address the following questions: 1) Is there genetic differentiation between anther-smut fungi parasitizing the closely related plant species *S. nutans* and *S. italica*? Are there hybrids and/or spill-overs in anther-smut fungi? 2) Is the population genetic structure of the anther-smut fungi parasitizing *S. nutans* and *S. italica* congruent with those of their hosts? 3) What is the divergence history of these anther-smut fungi on their hosts? Did genetic divergence occur with gene flow? Does the divergence history of the fungi mirror that of their hosts? In order to address these questions, we used genetic markers (nuclear microsatellite markers and/or chloroplastic SNPs) in both plants and anther-smut fungi to analyse diseased material (171 plant-pathogen pairs) collected across Western Europe and infer population genetic structures. We used approximate Bayesian computation (ABC) to compare genetic divergence scenarios in anther-smut fungi in order to assess whether the genetic divergence history in anther-smut fungi mirrored that inferred previously for their host plants. We also sequenced 53 genomes of *Microbotryum* fungi and analysed them together with 46 available *Microbotryum* genomes to assess whether the inferences on population genetic structure and gene flow based on microsatellite markers hold at the genome-wide levels and whether there were spill-overs or gene flow from other sympatric *Microbotryum* species than those analysed with microsatellite markers.

## Materials and Methods

### Fungal and plant materials

We analyzed 171 anther-smut fungi collected from diseased plant individuals of *S. nutans* and *S. italica* that were sampled in 55 distinct geographic sites across Europe, including the United Kingdom, France, Belgium, Switzerland, Italy, Germany and Norway (Table S1, Fig. 1A). Diseased plant individuals were stored in individual paper envelops kept in plastic bags filled with silica gel, in dark conditions at 8°C. Small pieces of leaves were used as raw material for DNA extraction of host plant individuals. Diploid spores of *Microbotryum* anther-smut fungi were collected in buds of infected flowers and grown as yeasts on potato dextrose agar (PDA) medium, and then stored at -20°C. Spores did not grow for a few samples older than two years. For DNA extraction of these strains, we used the dried spores directly collected from anthers of diseased flowers. In most cases, spores from anthers of a single flower were used and therefore corresponded to a single *Microbotryum* genotype (López-Villavicencio et al., 2007). When spore material was limited, we used multiple anthers from several flowers of the same plant. We never observed more than two alleles per strain and found low levels of heterozygosity per strain, indicating that we unlikely genotyped more than one fungal genotype.

**Figure 1:**
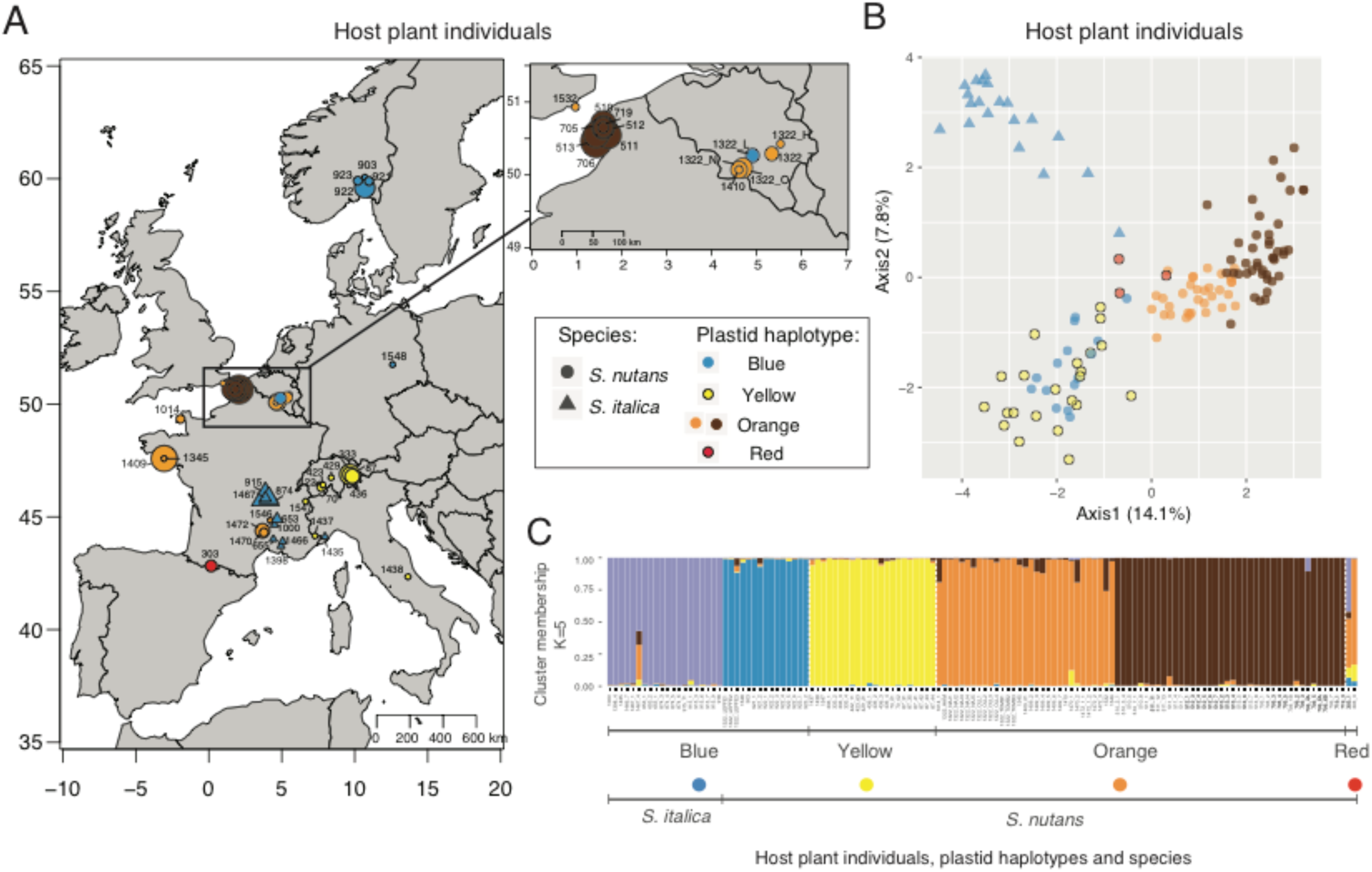
Population genetic structure of *Silene nutans* and *S. italica* host plants based on six plastid (chloroplast) SNPs and 21 microsatellite markers. A. Plastid haplotypes according to their geographic distribution. On the right, a zoom of the map in the northeastern region of France and Belgium is shown. The size of the symbols is proportional to the number of individuals sampled by site (1-9 plants). B. Principal component analysis (PCA), with information of plastid haplotypes. The first and second principal component axes are shown and the percentage of variance explained by each axis is indicated into brackets. C. Results of STRUCTURE on both host plant species for K=5 clusters. On panels A and B, symbol shape indicates species and color indicates plastid haplotypes. The two genetic clusters identified within the orange plastid haplotype based on the STRUCTURE analysis (panel C) are shown with light and dark orange colors, respectively.

**Table 1:**
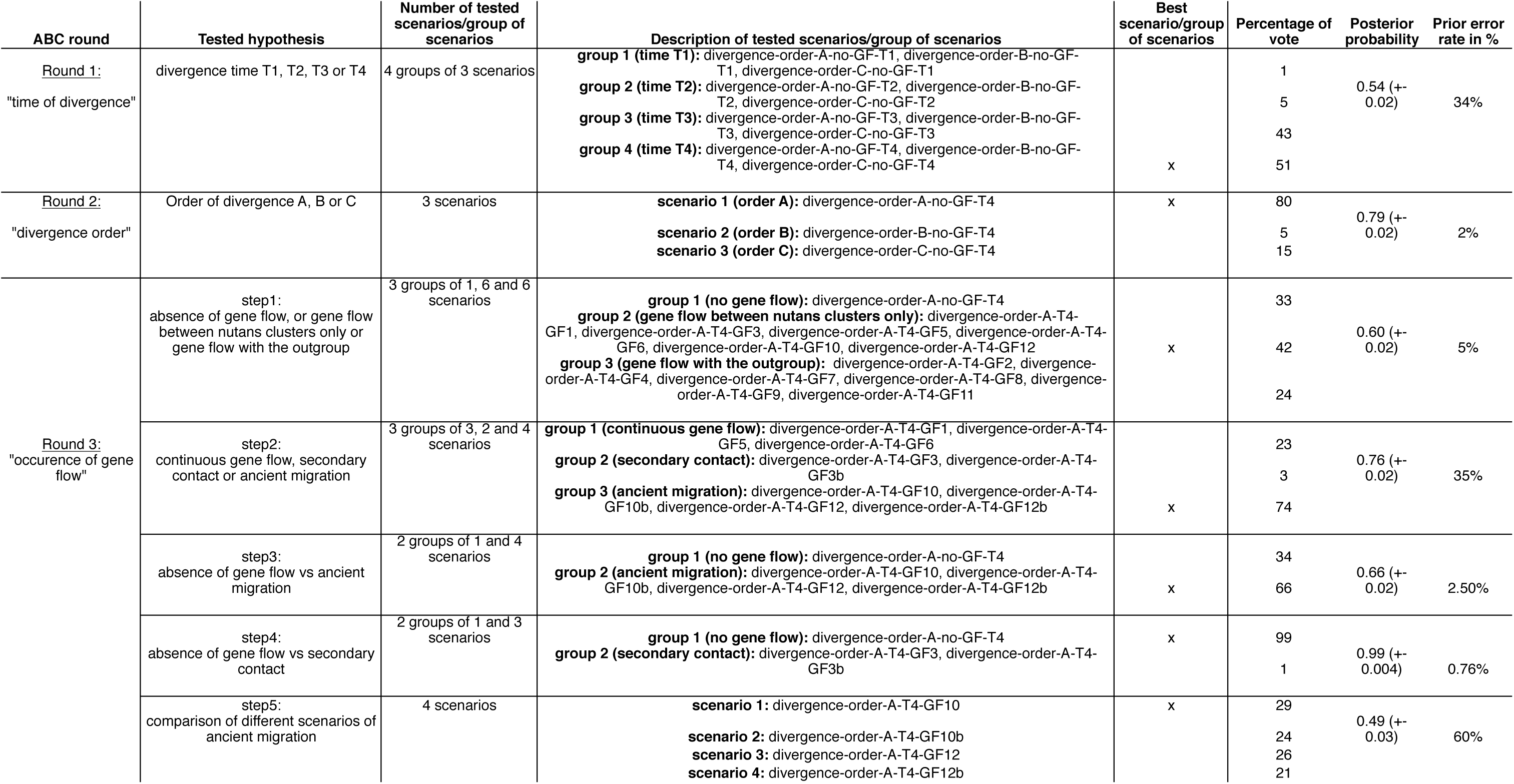
Comparisons of scenarios and group of scenarios used for approximate Bayesian computation (ABC) to reconstruct the fungi divergence history. Scenarios are described in Table S5 and Fig S5. Results of ABC random forest procedure are shown, with the percentage of vote for each scenario/group of scenarios, the posterior probability and the prior error rate.

### Genotyping and identification of species and genetic lineages of the *Silene* plants based on nuclear microsatellite markers and chloroplastic SNPs

We extracted plant DNA of at least one diseased plant individual per site using the NucleoSpin® 96 Plant II kit (Macherey-Nagel, Germany). We obtained DNA for 134 out of the 171 diseased plant individuals. At least one host individual was genotyped per site, except for six sites due to the lack of plant material (Table S1). In addition, we genotyped three plant individuals for which no anther-smut fungus was isolated but strains were isolated at the same sites (Table S1). Using morphological criteria of the host plant species, sample collectors initially identified 153 *S. nutans* and 18 *S. italica* diseased individuals. However, as the morphologies of *S. nutans* and *S. italica* are very similar, misidentification could occur. Therefore, we sequenced four chloroplast fragments (*psbA*, *LF*, *MATK*, *GS*; Lahiani et al., 2013) to check species identity of the diseased plants collected in southeastern France and in Italy, where both *S. nutans* and *S. italica* can be found. We thus reassigned one individual to *S. nutans* and seven individuals to *S. italica,* while one host individual could not be assigned to either *S. nutans* or *S. italica* species (from the pair # 1436). We excluded this host-pathogen individual pair from our analyses. We thus had 145 diseased *S. nutans* plants and 25 diseased *S. italica* plant (Table S1).

To further genotype the plant individuals for studying the population genetic structure, we used a combination of plastid (chloroplast) SNPs and nuclear microsatellite markers as previously described (Godé et al., 2014; Martin et al., 2016; Martin et al., 2017). For plastid markers, using the KASPAR® protocol we genotyped six SNPs, named Cp42, Cp397, Cp540, Cp656, Cp730 and Cp804, and polymorphic for [T/G], [A/C], [C/T], [G/T], [C/T], and [T/G], respectively. Individual haplotypes were defined as combinations of allelic states for all six SNPs. For nuclear markers, we used six multiplexes genotyping 24 microsatellite markers (Table S2). We followed the previously published protocols (Godé et al., 2014; Martin et al., 2016; Martin et al., 2017), except that we used different dye colors, bought from Eurofins Genomics. We used the Multiplex PCR Kit (Qiagen) following manufacturer instructions for PCR reactions, performed separately for each multiplex in 15 µL volume containing 3 µL of DNA, 3.4 µL of H2O, 7.1 µL of multiplex

PCR Kit (Buffer 2X, Qiagen, USA), and 1.4 µL of the primer mix. The primer mix included 2 µM of unlabelled forward and reverse primers and 0.5 µM or 0.75 µM of the labelled forward primer depending on the dye label. We used the same PCR cycling program as described in (Godé et al., 2014) with some modifications of the final elongation for multiplex 1 (Table S3A-C). We checked successful PCR amplifications on 2% (w/v) agarose gel electrophoresis. We outsourced genotyping at the Gentyane Genotyping Platform (INRA, Clermont, France) and scored alleles with GENEMAPPER v.4.0 (Applied Biosystems). We excluded three markers (SIL18, SIL26, SIL42) for which we had less than 50% of the individuals successfully genotyped. Given the material available, we could genotype 136 host plant individuals using both plastid SNPs and nuclear microsatellite markers. All 136 genotyped host individuals had a determined plastid haplotype and a determined genotype for at least 50% of the 21 remaining nuclear microsatellite markers. The genotypes are deposited on Dryad (accession available upon acceptance).

### Genotyping and species identification of the *Microbotryum* fungal strains: microsatellite markers and ITS sequences

We extracted fungal DNA using the Chelex protocol (Biorad, USA) following (Giraud, 2004). We extracted DNA from the 170 strains of anther-smut fungi, either from mixes, stored at -20°C, of haploid sporidia resulting from clonal growth after meiosis or from diploid teliospores directly collected in anthers of diseased flowers and stored at 10°C. Diploid genomes were therefore genotyped. DNA was diluted half-fold for PCR amplification. We used the internal transcribed spacer (ITS) to check that the genus of the fungal strains belonged to the *Microbotryum* genus. To genotype the fungal strains, we used 22 microsatellite markers arranged into multiplex (Table S4 ; (Fortuna et al., 2016; Giraud et al., 2008)). We used the Multiplex PCR Kit (Qiagen) following manufacturer instructions for PCR reactions which were performed separately for each multiplex as described above. We used the PCR cycling programs as in previous studies (Fortuna et al., 2016; Giraud et al., 2008; Table S3C-D). We checked the success of PCR amplifications on 2 % agarose gel electrophoresis. For genotyping, we pooled multiplexes 7 and 8 as a single multiplex. We outsourced genotyping at the Gentyane Genotyping Platform (INRA, Clermont, France) and scored alleles with GENEMAPPER v.4.0 (Applied Biosystems). We could determine the genotypes for all strains for at least 50% of the 22 markers. We identified 21 fungal strains that were likely siblings of other fungal strains based on null genetic distances and we excluded them from the STRUCTURE software analyses (see below).

### Population genetic structure based on microsatellite markers

To analyze and compare host and pathogen population genetic structures, we used a combination of three complementary approaches using microsatellite nuclear markers on both the *Silene* host plants and anther-smut fungal datasets. First, we used the model-based Bayesian clustering approach implemented in the software STRUCTURE version 2.3.4 (Pritchard, Stephens, & Donnelly, 2000). The program performs partitions of multilocus genotypes into genetic clusters and assigns individuals to genetic clusters, minimizing the departure from expected frequencies and linkage equilibrium among loci. We tested an admixture model with correlated frequencies and no prior information for K = 2 to K = 10 clusters. A total of 10 repetitions were run for each K value. We used 50,000 samples as a burn-in period and 100,000 samples per run for the Monte Carlo Markov Chain (MCMC) replicates. Cluster assignment probabilities were computed using the CLUMPP program (Jakobsson & Rosenberg, 2007) implemented in the R package {Pophelper}. We used the R package {Pophelper} (https://github.com/royfrancis/pophelper) to build the barplots. We choose as the biologically most relevant K value the finer population structure, as the highest K value for which a new cluster could be identified with individuals highly assigned to it, the new cluster at {K+1} having only admixed individuals (i.e. mean membership coefficient <0.80 to the given cluster). For the anther-smut fungus dataset, genetic data were haploidized as individuals were highly homozygous. High homozygosity levels might bias inferences in the software STRUCTURE as these are based on Hardy-Weinberg expectations in a diploid setting (Pritchard et al., 2000). Furthermore, 21 strains were removed, appearing as siblings of other strains, with identical genotypes. *Microbotryum* fungi have one obligatory sexual event before plant infection, so that clonemates cannot be found in different plants, but high selfing rates may allow the same genotype to be found in neighbor plants if they are parasitized by the same diploid spores or offspring (Giraud, 2004). For the host plant dataset, genetic data were kept as diploid and five individuals were removed as they had identical genotypes as other neighbor plants and may be clonemates. We identified clonemates and siblings in both datasets using the dist(X) function in the R package {Ape} (Paradis, Claude, & Strimmer, 2004) and considered two individuals to be siblings if their distance was equal to 0. We then used two methods for assessing population genetic structure that do not assume outcrossing or a lack of linkage disequilibrium. We performed a discriminant analysis of principal components (DAPC) using the R package {ADEGENET} (Jombart, 2008; Jombart & Ahmed, 2011) and used a principal component analysis (PCA) with the dudi.pca function using the R package {ade4} (Dray & Dufour, 2007) on the entire set of individuals in both datasets. Maps showing genotypes per locality were drawn using the R package {maps} (Becker, Wilks, Brownrigg, Minka, & Deckmyn, 2017) and {mapplots} (Gerritsen, 2013). Scatter plots were performed using the R package {ggplot2} (Wickham, 2009).

### Population statistics of genetic diversity and structure based on microsatellite markers

We computed, using the R package {diveRsity} (Keenan, McGinnity, Cross, Crozier, & Prodöhl, 2016), the following estimates of genetic diversity per locus, site, genetic cluster identified in the STRUCTURE analysis and/or species for the host plants and for the anther-smut fungi datasets: the number of alleles, allelic richness (A_R_), observed heterozygosity (H_O_), expected heterozygosity (H_E_), the fixation indexes (F_ST_, F_IS_ and F_IT_), and the Jost’s D statistics corresponding to the fraction of allelic variation found among genetic clusters. We tested whether genotype frequencies fitted the Hardy-Weinberg expectations using a standard Chi² goodness of fit method and assessed the significance of F_ST_, F_IS_ and F_IT_ values using a bootstrap procedure with 1000 iterations and calculating 95% confidence intervals. Sites with less than three sampled individuals were pooled with other sites of the same genetic lineage when they were closer than 1/10th of latitude or longitude or were otherwise excluded from the analyses. We excluded nine sites for the plants (site # 1014, 303, 1532, 719, 1546, 1547, 1437, 429, 1548, 1438) and 13 sites for the fungal pathogens (site # 1068, 1014, 303, 1532, 719, 1546, 940, 1547, 1437, 333, 1249, 6809, 1548). In total, we considered 20 groups of sites for the host and 19 groups of sites for the pathogen. We included sibling individuals but excluded individuals with admixed membership between genetic clusters inferred from the STRUCTURE analysis (i.e. individuals with mean membership coefficient <0.80 to the given cluster), which may be due to low assignment power or admixture. To take into account differences of sample size between sites, we also estimated allelic richness using ADZE (Szpiech, Jakobsson, & Rosenberg, 2008) which corrects for sample size difference. Calculations were performed using a standardized sample size of N=3, corresponding to the smallest number of observations per site. We tested for mean differences in diversity statistics between species and genetic clusters using a Wilcoxon rank sum test and a Kruskal-Wallis rank sum test in the R software v3.5.3, respectively, considering values across groups of sites. To study isolation-by-distance patterns in the *S. nutans* and fungi parasitizing *S. nutans* datasets, we computed correlations between matrices of genetic and geographic distances of plant and fungal populations using a Mantel test. Genetic distances between populations at each group of sites were calculated as the Nei’s distance (Nei, 1972) using the dist.genpop() function of the R package {ADEGENET} (Jombart, 2008; Jombart & Ahmed, 2011). Geographic distances were calculated using the distm() function of the R package{geosphere}. We computed Mantel tests using the mantel.rtest() function of the R package {ade4} with 1,000,000 resamples for the null distribution. We also studied the correlations between matrices of genetic distances of plant and fungal populations using a Mantel test as above. To remove the effect of the correlation between genetic and geographic distances in the correlation between matrices of genetic distances, we also performed a partial Mantel test using the partial.mantel.test() function of the R package {ncf} with 1,000,000 resamples for the null distribution (Feurtey et al., 2016). We considered 16 group of sites, the same both for the host and the pathogen, corresponding to *S. nutans* host plants and fungal strains belonging to the genetic cluster parasitizing S*. nutans.* We excluded sites of *S. italica* host plants and fungal strains belonging to the genetic cluster parasitizing *S. italica,* and one site studied in the host but not studied in the pathogen (site # 333). We investigated the occurrence of recent events of effective population size reduction (i.e. bottlenecks) within the identified fungal genetic clusters using the program BOTTLENECK version 1.2.02 (Piry, Luikart, & Cornuet, 1999).

### Inference of anther-smut fungi divergence history based on microsatellite markers

We studied the divergence history of the anther-smut fungi sampled on the *S. nutans* lineage complex using an approximate Bayesian computation-random forest (ABC-RF) procedure that performs ABC inferences based on the machine learning tool named “random forest” (Breiman, 2001; Pudlo et al., 2016; Raynal et al., 2019). The divergence history of the host had already been previously inferred (Martin et al., 2016; Martin et al., 2017; Van Rossum et al., 2018). To increase our power to disentangle between different evolutionary scenarios, we performed model choice and parameter estimation by comparing scenarios or groups of scenarios in sequential rounds, each round testing a particular type of evolutionary event, either divergence time, order of divergence or presence of gene flow (Estoup, Raynal, Verdu, & Marin, 2018; Liu et al., 2019); Table 1; Table S5). We built scenarios based on the genetic clusters obtained using microsatellite markers and on previous analyses of divergence of anther-smut fungi and their host species (Badouin et al., 2017; Gladieux et al., 2013; Martin et al., 2016; Martin et al., 2017; Van Rossum et al., 2018). The tested scenarios varied regarding the time of divergence, the relative order of divergence of fungal genetic clusters and the occurrence of gene flow among genetic fungal clusters (Table S5). We did not include scenarios with variation in effective population sizes as the BOTTLENECK analysis did not identify any signature of effective population size reduction (see ‘Results’ section). To test if the order of sequential rounds had an effect on the outcome of the analysis, we tested the time of divergence and the relative order of divergence as either the first or the second round (Note S1). We used as populations the fungal genetic clusters identified through population structure analyses and removed fungal strains that were likely siblings of other fungal strains in each genetic cluster.

We ran the ABC procedure modified from (Liu et al., 2019). Briefly, we simulated datasets with 22 fungal microsatellite markers using the ABCtoolbox program (Wegmann, Leuenberger, Neuenschwander, & Excoffier, 2010) and used fastsimcoal 2.5 for coalescent-based simulations (Excoffier & Foll, 2011). We simulated 10,000 genetic datasets per scenario using coalescent simulations with model parameters drawn from prior distributions (Table S6A). We set prior distributions based on previous analyses of divergence of anther-smut fungi and of their host species (Badouin et al., 2017; Branco et al., 2018; Gladieux et al., 2013; Martin et al., 2016; Martin et al., 2017; Van Rossum et al., 2018). We set a generation time of one year for the pathogen based on its life cycle (Thrall, Biere, & Antonovics, 1993). We estimated the following parameters: effective size of each genetic cluster (*N*), divergence time and migration rate per generation between two genetic clusters *x* and *y* (*Txy* and *mxy*, respectively). We computed 16 summary statistics for the observed and simulated datasets with the program arlsumstats (Table S6B; (Excoffier & Lischer, 2010). We assumed a generalized stepwise model of microsatellite evolution (Estoup, Jarne, & Cornuet, 2002) and allowed the mutation rate to vary across microsatellite markers. We drew locus-specific mutation rates from a gamma distribution (α,α/μ) in which μ is the mutation rate per generation and α is a shape parameter (Cornille et al. 2012; Liu et al., 2019). We used the R package {abcrf} v1.7.0 (Pudlo et al., 2016) to compute a classification vote through ABC-RF, representing the number of times a scenario was selected among classification *n* trees of the constructed random forest. We chose n=500 trees. For each round, we selected the scenario or group of scenarios with the highest number of classification votes. We computed the posterior probability and the prior error rates over 10 replicated analyses. We performed a linear discriminant analysis (LDA) in the R package {abcrf} v1.7.0 (Pudlo et al., 2016) on the simulated and observed datasets to visually check the fit of the model to the observed data. Finally, we performed parameter inferences using the group of models eventually selected.

### Sequencing data and genome assemblies

We performed whole-genome sequencing for 23 anther-smut fungal strains parasitizing *S. italica* and 30 strains parasitizing *S. nutans*. Out of these 53 strains, 38 strains belonged to the 170 host-pathogen pairs that were genotyped with microsatellite markers (Table S7). Due to limited fungal material, we sequenced six additional strains that were collected at our studied sites but not genotyped for microsatellite markers and nine additional strains collected at three new sites (Table S7). We extracted DNA from fresh spores stored at -20°C using the Nucleospin Soil Kit (Macherey-Nagel, Germany). Genomes were sequenced using Illumina 150 bp paired-end sequencing technology at 46X coverage on average. We also included outgroups using available whole-genome sequences of 56 anther-smut fungal strains parasitizing closely-related host species of *S. nutans* and *S. italica* occurring in the studied geographic range (Le Gac, Hood, Fournier, et al., 2007; Refrégier et al., 2008): 33 *M. lychnidis-dioicae* strains parasitizing *S. latifolia* (Badouin et al., 2017), 19 *M. silenes-dioicae* strains parasitizing *S. dioica* (Badouin et al., 2017) and four *M. violaceum* var *paradoxa* strains parasitizing *S. paradoxa* (Branco et al., 2018) (Table S7). We downloaded raw data publicly available from the NCBI SRA under the BioProject IDs PRJNA295022 and PRJEB16741. We thus analysed a total of 109 *Microbotryum* genomes. For read mapping, we used as reference genome the high-quality haploid genome assembly of the MvSn-1249 *M. violaceum s. str.* strain corresponding to the a_2_ mating type (collected on *S. nutans*) previously obtained with P6/C4 Pacific Biosciences SMRT technology and annotated for gene models (Branco et al., 2017). The MvSn-1249-A2 assembly was accessed from GenBank BioProject accession number PRJEB12080 (BioSample ID: SAMEA3706514, assembly: GCA_900014965).

### Genome read mapping and variant calling procedure

We performed read mapping and SNP calling of the 109 analysed genomes against the MvSn-1249-A2 genome as previously described (Branco et al., 2018; Hartmann, Rodríguez de la Vega, Brandenburg, Carpentier, & Giraud, 2018). First, we trimmed Illumina raw reads for sequence quality and removed adapter sequences using the software Cutadapt v1.8.3 (Martin, 2011) with the options: -q 10, 10; –minimum-length 50; -a AGATCGGAAGAGCACACGTCTGAACTCCAGTCAC; - AAGATCGGAAGAGCGTCGTGTAGGGAAAGAGTGTAGATCTCGGTGGTCGCCGTATT.

We aligned trimmed reads using the short read aligner bowtie2 v2.1.0 (Langmead, Trapnell, Pop, & Salzberg, 2009) with the three following software options: –very-sensitive-local; –phred33; -X 1000. We removed PCR duplicates using the MarkDuplicates tool of Picard tools version 2.14.1 (http://broadinstitute.github.io/picard) and performed local realignment of mapped reads using the tools RealignerTargetCreator and IndelRealigner of the Genome Analysis Toolkit (GATK) version 3.8 (McKenna et al., 2010) to improve alignment accuracy in indel regions. Mean alignment rates to the reference genome ranged from 53 to 96% (Table S7). For SNP calling, we used GATK version 3.7 (McKenna et al., 2010) and ran HaplotypeCaller on each strain individually using a diploid mode. Then, we used GenotypeGVCFs on a merged gvcf variant file to perform joint variant calls. We filtered SNP calls for quality using VariantFiltration and following the GATK good practices for hard-filtering of variants (QUAL < 250; QD < 2; MQ < 30.0; -10.5 > MQRankSum > 10.5; -5 > ReadPosRankSum > 5; FS > 60; SOR > 3) and masked SNPs for repeats. We found 41 SNPs in the strain MvSn-1249-A2 (the very same fungal strain as the reference genome also sequenced with Illumina), indicating low rates of false positives in our SNP calling. We only kept bi-allelic SNPs with a high genotyping rate (> 90%) on autosomes (i.e., excluding SNPs on the mating-type contigs MvSn-1249-A2-R1_MC15 and MvSn-1249-A2-R1_MC16). We excluded SNPs on the mating-type chromosomes for population genomics analyses as they exhibit suppressed recombination on 90% of their lengths (Branco et al., 2018). Removing the strain MvSn-1249-A2, we kept 108 fungal strains and 1,305,369 SNPs for population genomic analyses.

### Population genomics analyses

To analyze the population genetic structures of anther-smut fungi parasitizing several *Silene* species using the genome-wide SNP dataset called on autosomes, we used the model-based Bayesian clustering approach implemented in the software STRUCTURE version 2.3.4 (Pritchard et al., 2000), from K=2 to K=10 clusters, and performed a PCA, similarly as for the microsatellite datasets. For the STRUCTURE analysis, we selected a set of 233 unlinked SNPs randomly distributed at intervals of 100 kb along the autosomes, linkage disequilibrium decaying over 50-100 kb in *Microbotryum* populations (Badouin et al., 2017). We used the R package {Pophelper} (https://github.com/royfrancis/pophelper) to perform barplots. We performed the PCA on all SNPs (but excluding missing data and heterozygote genotypes) using the --pca command of the Plink v1.9 software (Chang et al., 2015; Purcell et al., 2007). To study footprints of gene flow between fungal strains, we performed a phylogenetic network analysis implemented in splitsTree (Huson, 1998; Huson & Bryant, 2006) with the neighbornet method.

## Results

### Population genetic structure and diversity of *Silene nutans* and *S. italica* host plants

After checking plant species identity with molecular markers, we retained 145 diseased *S. nutans* plants and 25 diseased *S. italica* plants (Table S1; Fig 1A). The 21 plant nuclear microsatellite markers distinguished well *S. nutans* and *S. italica* both in STRUCTURE analyses from K=2 and in the principal component analysis (PCA; Fig 1B-C; Fig S1).

Within *S. nutans*, we identified four distinct haplotypes based on the six plastid SNPs, retrieving in our sample the four major genetic lineages of *S. nutans* previously identified in Western Europe (Fig 1A; (Martin et al., 2016; Martin et al., 2017; Van Rossum et al., 2018). The blue, yellow, orange, and red haplotypes were found, as in previous studies, in the northeastern, southeastern, western and southwestern parts of Europe, respectively. All *S. italica* samples had a blue plastid haplotype (Fig 1A). Within *S. nutans,* the PCA based on the 21 plant nuclear microsatellite markers mainly differentiated the orange genetic lineage from the other ones (Fig 1B; Fig S1B) and the STRUCTURE analysis detected four genetic clusters, corresponding to the different plastid haplotypes, except for the red haplotype for which we had too few samples (Fig 1C; Fig S1A). The STRUCTURE analysis further identified two well-separated genetic clusters within the *S. nutans* orange plastid haplotype (Fig. 1C). These two genetic clusters also appeared separated on the second axis of the PCA and corresponded to plant individuals collected in the Northern part of France *versus* Belgium and Western part of France (Fig S1), in agreement with previous findings (Martin et al., 2016). From the STRUCTURE analyses, we identified five host individuals with admixed cluster memberships (mean membership coefficient <0.80 to the given cluster), which may be due to low assignment power or admixture between genetic clusters. We found no further clear subdivision within *S. nutans* at higher K values and no clear population subdivision within *S. italica* (Fig S1A).

We found no significant differences in allelic richness neither between *S. nutans* and *S. italica* (Wilcoxon rank sum (WRS) test on 20 groups of sites, W = 12, p-value = 0.169; Table S8A) nor between the four *S. nutans* genetic clusters (Kruskal-Wallis rank sum (KWRS) test on 16 groups of sites, Chi² = 4.92, degree of freedom (df) = 3, p-value = 0.178; Table S8A). In *S. nutans,* levels of observed heterozygosity were significantly lower than those of expected heterozygosity under Hardy-Weinberg equilibrium for most loci, the mean F_IS_ value per marker being 0.33 (Table S2). In *S. italica,* levels of observed and expected heterozygosities were not significantly different for most markers (78%; Table S2), the mean F_IS_ value per locus being 0.37 (Table S2). As expected, pairwise F_ST_ and Jost’s D indicated higher divergence between *S. italica* and *S. nutans* (mean F_ST_=0.32; mean Jost’s D=0.57; Table S9A) than between genetic clusters of *S. nutans* (mean F_ST_=0.20; mean Jost’s D=0.35; Table S9A; KWRS test for both statistics, W = 0, p-value = 0.010). For *S. nutans,* we found on average lower pairwise F_ST_ and Jost’s D values between the yellow Western cluster and the two orange Western and Northwestern genetic clusters (mean F_ST_=0.17; mean Jost’s D=0.28; Table S9A) than between the blue Eastern genetic cluster and the Western genetic clusters (mean F_ST_=0.24; mean Jost’s D=0.42; Table S9A), although the difference was not significant (KWRS test for F_ST_, W = 8, p-value = 0.200; KWRS test for Jost’s D, W = 9, p-value = 0.100). The correlation between the matrices of pairwise genetic and geographic distances for *S. nutans* (tested on 16 sites) was significantly positive (Mantel test: r = 0.54, p-value < 0.001), indicating an isolation-by-distance pattern.

### Differentiation between anther-smut fungi parasitizing *Silene nutans* and *S. italica*

Population genetic structure analyses separated *Microbotryum* strains into two main genetic clusters, corresponding to their host plant of sampling, *S. italica* and *S. nutans*. The host of collection separated strains according to the first axis of the PCA (Fig 2B). The STRUCTURE analysis and the discriminant analysis of principal component (DAPC) also delimitated two clear fungal genetic clusters according to the host of sampling at K=2 (Fig 2C; Fig S2). In the PCA (Fig. 2B), four *Microbotryum* strains sampled on *S. nutans* clustered with strains parasitizing *S. italica* and one *Microbotryum* strain sampled on *S. italica* clustered with strains parasitizing *S. nutans*. These strains may correspond to spill-overs, i.e. cross-species disease transmissions, between the two studied *Silene* species or from other *Silene* species. We excluded for further analyses the 11 strains that had their main cluster memberships < 80% in the STRUCTURE analysis at K=2 (Fig 2C), which can be due to low assignment power or admixture.

**Figure 2:**
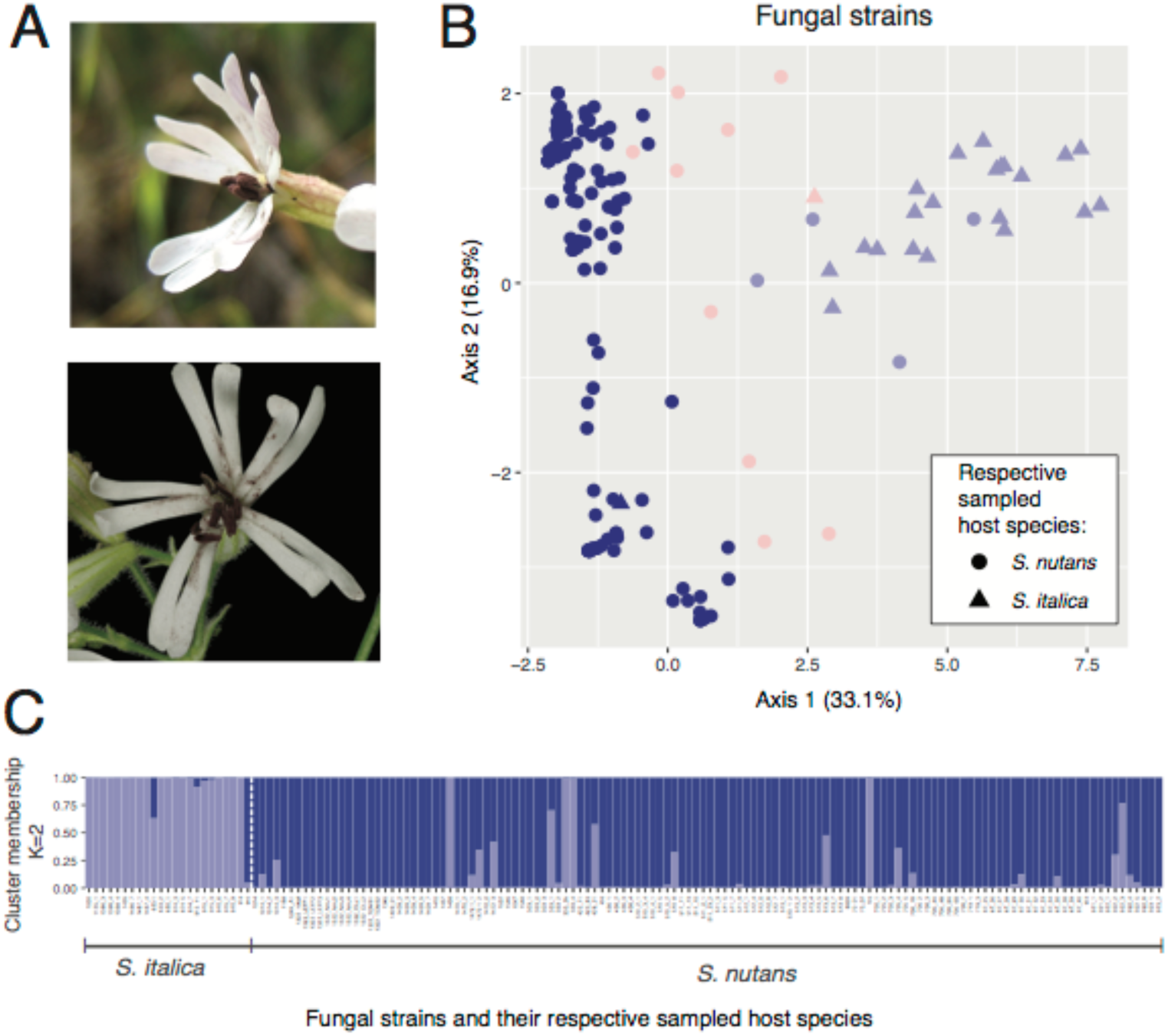
Population genetic structure of anther-smut fungi (*Microbotryum*) parasitizing *Silene nutans* and *S. italica* based on 22 microsatellite markers. A. Anther-smut fungi parasitizing *S. italica* (top; photo credit M.E. Hood) and *S. nutans* (bottom; photo credit M. Strack van Schijndel). B. Principal component analysis (PCA). The first and second principal component axes are shown and the percentage of variance explained by each axis is indicated into brackets. The symbol shape indicates the sampling host species. The dark and light blue colors indicate assignment probability >0.80 to the two corresponding clusters identified in the STRUCTURE analysis for K=2 (see panel C). Pink color indicates strains for which no cluster could be assigned with a probability >0.80. Genetic variance explained by the principal component axes is shown in the right bottom corner. C. STRUCTURE barplot for K=2. Strains are ordered according to their sampling host species.

Anther-smut fungi parasitizing *S. italica* had on average higher levels of allelic richness (WRS test on 19 groups of sites, W = 48, p-value = 0.004) and observed heterozygosity (WRS test on 19 groups of sites, W = 30, p-value = 0.014) than anther-smut fungi parasitizing *S. nutans* (Table S8B). Levels of observed heterozygosity were significantly lower than heterozygosity expected under Hardy-Weinberg equilibrium for eight markers for *Microbotryum* fungi parasitizing *S. nutans* and six markers for *Microbotryum* fungi parasitizing *S. italica* (Table S4).

### Similar genetic structure in anther-smut fungi as in their *Silene nutans* and *S. italica* host plants

The existence of additional levels of population genetic structure within fungal strains was indicated by the STRUCTURE barplots (Fig S2), the second and third principal components of the PCA that explained 27.9 % of the total variance between strains (Fig 2B). To investigate the population structure within each of the two main genetic clusters of anther-smut fungi, corresponding mainly to populations parasitizing *S. nutans* and *S. italica*, respectively, we analyzed them separately, which confirmed the existence of a strong population structure within anther-smut fungi parasitizing *S. nutans* (Fig 3A). The first two PCA axes separated fungal strains into three distinct genetic clusters, corresponding to the three most frequent plastid haplotypes identified in the host plant (Fig 3A), which was also supported by the DAPC and the STRUCTURE analyses (Fig 3B; Fig S3). We named these three fungal genetic clusters the blue Eastern, yellow Western and orange Western *Microbotryum* clusters, in reference to their *S. nutans* host plastid haplotypes. In contrast to the host plant *S. nutans*, we found no clear subdivision in the orange Western anther-smut fungal cluster (Fig S3). We found no genetic clusters specific to strains sampled from a host with a red haplotype, perhaps due to the low strain number. We excluded for further analyses the two admixed fungal strains (cluster memberships <0.80 in a given cluster) and the three fungal strains sampled on a host with a red haplotype. We found no further population structure in the set of 28 anther-smut strains parasitizing *S. italica* (Fig S4), as for the host plant.

**Figure 3:**
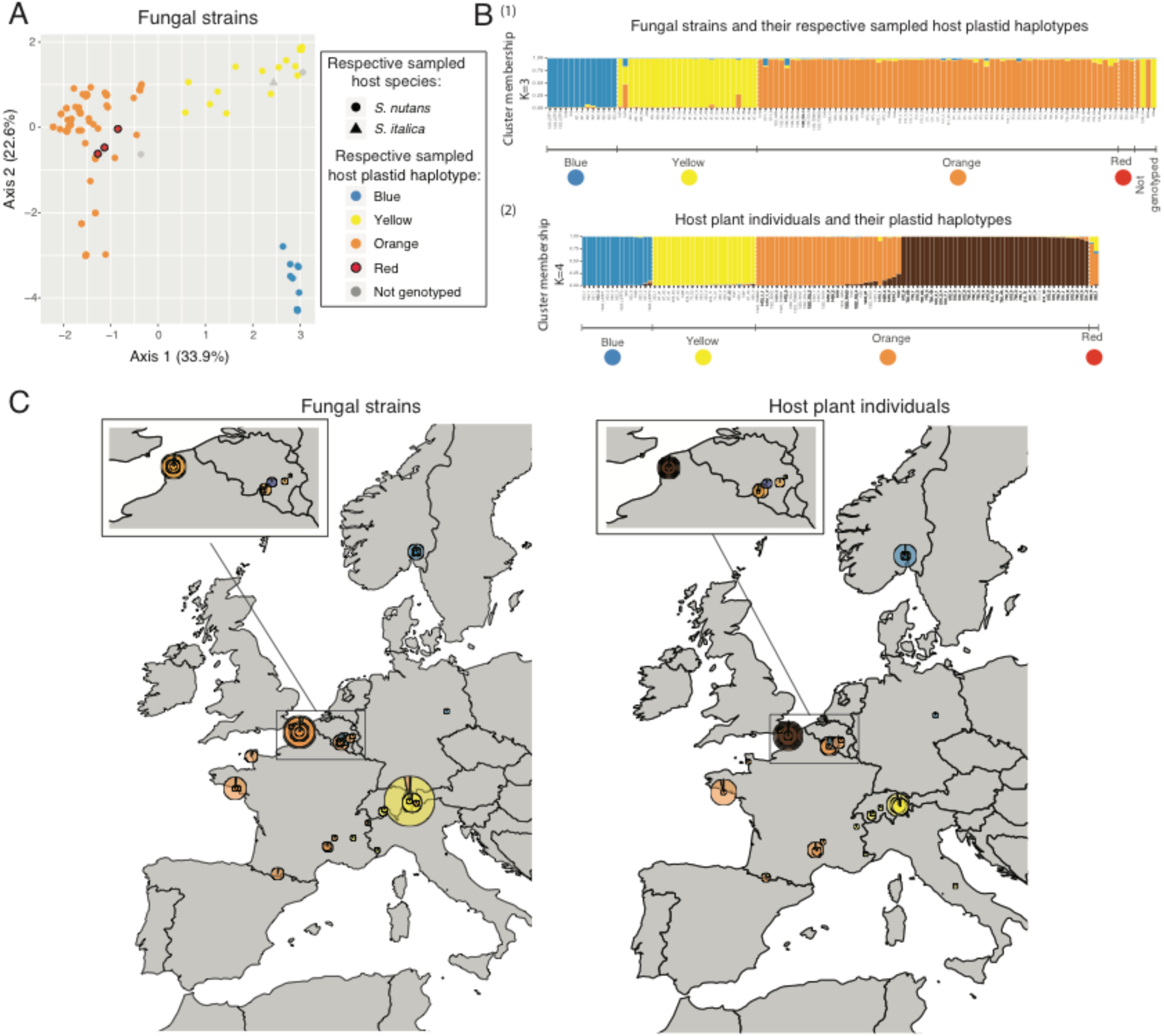
Congruence of population genetic structure between *Silene nutans* host plants and their anther-smut fungal (*Microbotryum*) strains based on microsatellite markers. A. Principal component analysis (PCA) on the fungal strains with information of the population genetic structure of their hosts. The first and second principal component axes are shown. Symbol shape indicates the sampling host species and color indicates the plastid haplotypes of the sampling host. B. STRUCTURE barplots for (1) K=3 in 112 *Microbotryum* strains and (2) K=4 in 110 *S. nutans* plant individuals. Fungal strains are ordered according to their sampling host plastid haplotypes. Host plant individuals are ordered according to their plastid haplotypes. C. Map of mean cluster membership proportions per site for (1) the 112 fungal strains in the STRUCTURE analysis at K=3 and (2) the 110 *S. nutans* plant individuals in the STRUCTURE analysis at K=4. The pie diameter reflects the sample size in the corresponding site (1 - 14 individuals).

All three fungal genetic clusters parasitizing *S. nutans* had similar levels of allelic richness (KWRS test on 16 groups of sites, Chi² = 1.33, df = 2, p-value = 0.514; Table S8B) and observed heterozygosity (KWRS test, Chi² = 0.67, df = 2, p-value = 0.716; Table S8B). The F_IS_ varied between genetic clusters from -0.11 to 0.82 (Table S8B). We found no significant differences in differentiation between the three fungal genetic clusters (KWRS tests for F_ST_ and Jost’s D, Chi² = 2, df = 2, p-value = 0.368; Table S9B). The significant correlation between the matrices of genetic and geographic distances between pairs of fungal populations parasitizing *S. nutans* (16 sites) indicated an IBD pattern (Mantel test: r = 0.62, p-value < 0.001). There was a significantly positive correlation between genetic distance matrices for the fungal and the *S. nutans* populations (16 sites; Mantel test: r = 0.80, p-value < 0.001), and it remained significant when controlling for the IBD effect (partial Mantel test: r = 0.70, p-value < 0.001). Altogether, our findings indicate strong congruence between the population genetic structures of the anther-smut fungi parasitizing *S. nutans* and its host plant *S. nutans* (Fig 3C). We found no signatures of recent reduction in effective population size in either of the three fungal genetic clusters parasitizing *S. nutans* nor the fungal genetic cluster parasitizing *S. italica* using the BOTTLENECK software (Piry et al., 1999) under either stepwise or two-phase models of microsatellite evolution (Table S10; one-tailed Wilcoxon signed rank test, all P-values>0.36).

### Inferred divergence history of anther-smut fungi on *Silene nutans* congruent with of their host plant

To study if the anther-smut fungi on *S. nutans* shared the same history of divergence into cryptic lineages as the one previously inferred for their host plants, the *S. nutans* cryptic species complex (Martin et al., 2016; Martin et al., 2017; Van Rossum et al., 2018), we used an approximate Bayesian computation random forest (ABC-RF) procedure. We compared various divergence scenarios for the three identified *Microbotryum* genetic clusters parasitizing *S. nutans.* We removed the 21 fungal strains that were likely siblings of other fungal strains, keeping 110 anther-smut fungal strains belonging to the blue Eastern, yellow Western and orange Western *Microbotryum* genetic clusters parasitizing *S. nutans* and we used as outgroup the 25 genotyped *Microbotryum* strains parasitizing *S. italica*.

We performed three rounds of scenario comparison, each testing a particular evolutionary event, including divergence times, divergence order and the occurrence of gene flow (Table 1; Table S5; Fig S5). At each round, we retained the inferred most likely evolutionary scenario to be used as backbone for the subsequent rounds. To assess our power to discriminate between scenarios, we checked posterior probability (Table 1) and assessed visually whether the observed data fell within the clouds of simulated data of the compared scenarios (Fig S6).

In the first round (“time of divergence”; Table 1, “round 1”), we tested four different time periods of divergence for various divergence orders (A, B or C; Fig S5A). The group of scenarios with strongest support included a first divergence occurring between 100,000-1,500,000 years ago, and a second divergence occurring between 0-20,000 years ago (posterior probability = 0.54 +-0.02, prior error rate = 34%; Table 1, “round 1”; Fig S6A). In the second round (“divergence order”; Table 1, “round 2”), we found with a high posterior probability for the scenario assuming a first divergence of the blue Eastern *Microbotryum* genetic cluster, followed by the divergence between the yellow and orange Western *Microbotryum* genetic clusters (posterior probability = 0.79 +-0.02, prior error rate = 2%; Table 1, “round 2”; Fig S6B). When testing the order of divergence between clusters as first round and the time of divergence as second round, we selected the same scenarios with a high posterior probability (Note S1; Table S11).

In the third round (“occurrence of gene flow”; Table 1, “round 3, step 1”), we compared in a first step three groups of scenarios differing in the clusters affected by gene flow. The most supported group of scenarios assumed gene flow among *Microbotryum* genetic clusters parasitizing *S. nutans* but not with *S. italica* (posterior probability = 0.60 +-0.02, prior error rate = 5%; Table 1, “round 3, step 1”; Fig S6C). In the following steps (Table 1, “round 3, steps 2 to 4”), we compared groups of scenarios of divergence with different timing of gene flow. Scenarios of divergence with ancient gene flow were the most supported, which suggested that episodes of ancient gene flow occurred during divergence or just following divergence of the three *Microbotryum* genetic clusters parasitizing *S. nutans* and then completely stopped (posterior probability = 0.76 +-0.02, prior error rate = 35%; Table 1, “round 3, step 2”; Fig S6D-F). However, we did not have enough power to differentiate between scenarios of ancient gene flow that involved different genetic clusters (either all three genetic clusters or only the two Western genetic clusters) or different times of past gene flow (either over a 0-20000 year period or a 50,000-500,000 year period; Table S5; posterior probability = 0.49 +-0.03, prior error rate = 60%; Table 1, “round 3, step 5”). We performed parameter inferences for the group of four models with ancient migration (“round 3, step 2”, group 3 see Table 1; Table S12). The inferred dates and order of divergence between fungal genetic clusters parasitizing *S. nutans* were consistent with the history of divergence previously inferred for the host plant *S. nutans* (Fig 4; Martin et al., 2016; Martin et al., 2017; Van Rossum et al., 2018).

**Figure 4:**
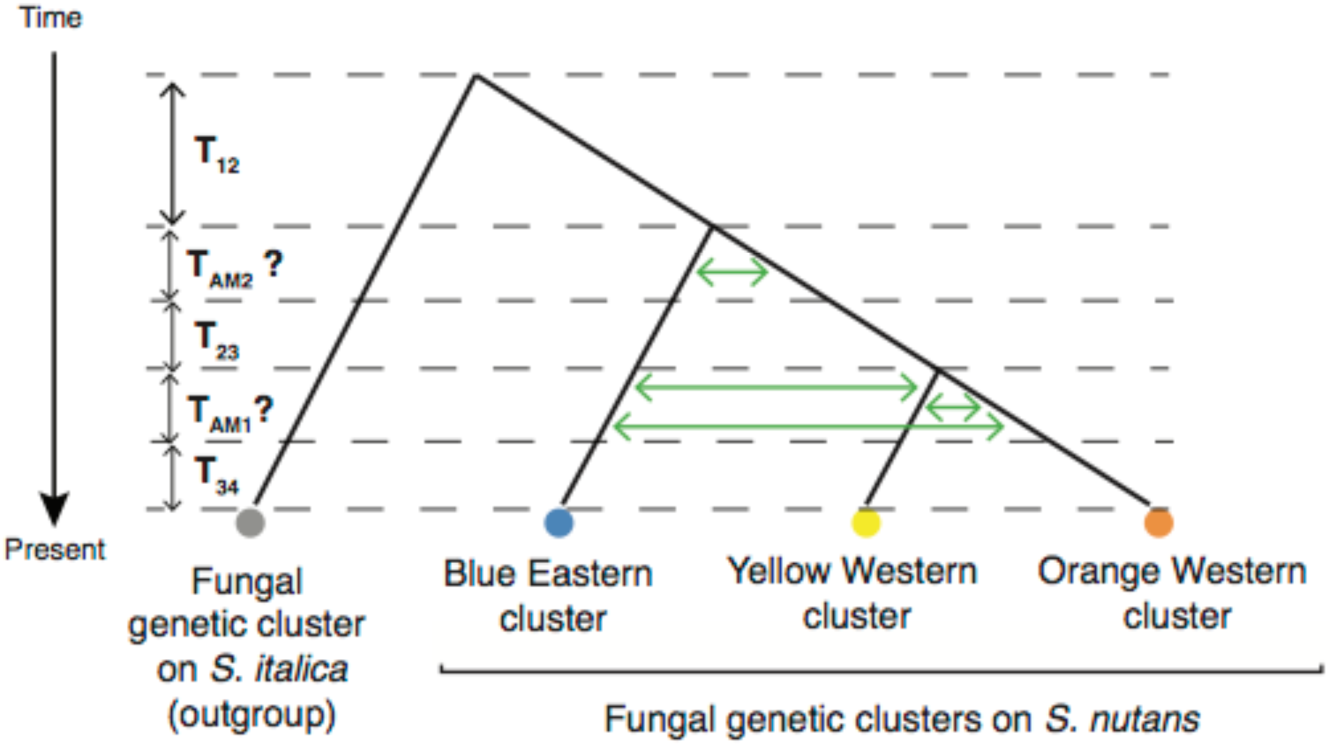
Most likely scenario for the divergence between the fungal genetic clusters inferred with approximate Bayesian computation (ABC).

### No genome-wide signatures of gene flow between anther-smut fungi on distinct *Silene* **species**

To check whether genome-wide data support the patterns inferred above on population genetic structure in the anther-smut fungi and lack of recent gene flow, we sequenced the genomes of 53 *Microbotryum* strains parasitizing *S. nutans* or *S. italica* (Table S7). We also used previously sequenced genomes of anther-smut fungi parasitizing species with overlapping geographical ranges in order to check if some of the strains that appeared as admixed or did not cluster according to their host species may actually correspond to other *Microbotryum* species. After filtration, we obtained 1,305,369 SNPs for 108 *Microbotryum* strains. We performed PCA analyses on the genome-wide SNPs and STRUCTURE analyses and confirmed strong population differentiation between fungal strains parasitizing *S. nutans* and *S. italica* (Fig 5A-B; Fig S7A). Only one strain parasitizing *S. nutans* clustered with strains parasitizing *S. italica,* similarly as found from microsatellite data (strain # 1438; Fig 2C), therefore likely being a genuine spill-over between *S. nutans* and *S. italica*. Two other strains parasitizing *S. nutans* clustered with strains parasitizing *S. latifolia.* These strains had admixed cluster membership (i.e. mean membership coefficient <0.80 to the given cluster) or clustered with strains parasitizing *S. italica* from microsatellite data (Fig 2C). These strains likely represent spill-overs between *S. nutans* and *S. latifolia*.

**Figure 5:**
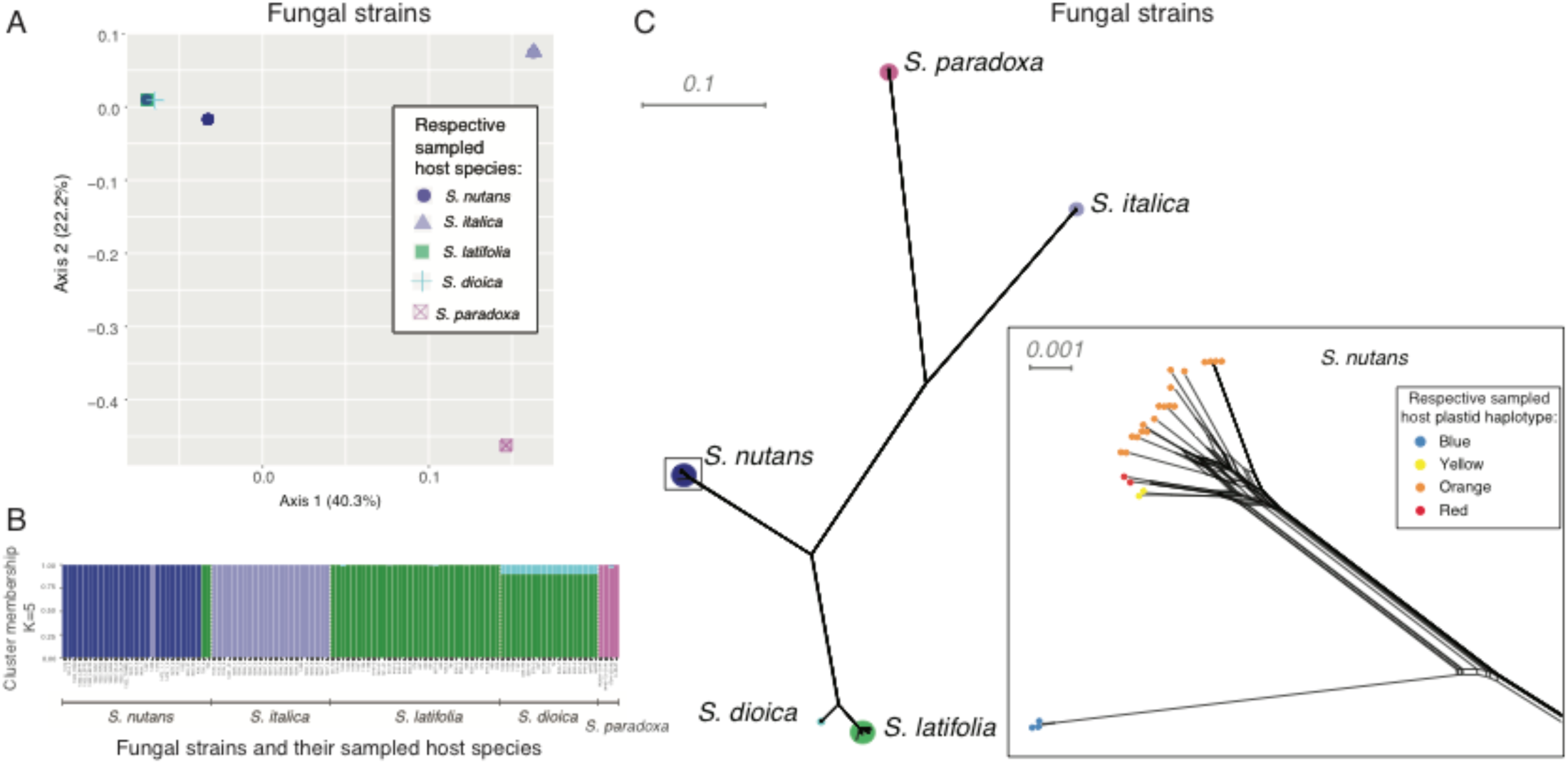
Absence of genome-wide signature of recent gene flow among anther-smut fungal strains parasitizing closely related *Silene* species. A. Principal component analysis (PCA) based on 1,305,369 genome-wide SNPs. B. STRUCTURE analyses based on 233 unlinked SNPs for K=5. The Y axis indicates the estimated membership proportions in the *K* clusters for each fungal strain (X axis). C. Neighbornet tree from a SplitsTree analysis based on 595,002 genome-wide SNPs with no missing data and heterozygote genotypes. The inner plot shows a zoom of the tip of the tree for strains parasitizing *S. nutans*. Information on the species or plastid haplotype of the host of sampling of the strains is given.

Genome sequences retrieved the same genetic structure as microsatellites for anther-smut fungi parasitizing *S. nutans,* with a strong differentiation between *Microbotryum* strains sampled on the *S. nutans* blue plastid haplotype on the one hand and those sampled on yellow, red or orange plastid haplotypes on the other hand (Fig S7B), and confirmed the absence of strong population structure within anther-smut fungi parasitizing *S. italica* (Fig S7C). We also found strong population differentiation between fungal strains parasitizing the three closely related host species of *S. nutans* and *S. italica* (Fig 5A-B). STRUCTURE analyses showed no signatures of recent admixture between strains parasitizing different hosts (Fig 5B).

The phylogenetic network analysis implemented in SplitsTree also supported the absence of recent gene flow at the genome-wide levels between the *Microbotryum* species parasitizing different hosts and the close genetic similarity of the three genetic clusters of anther-smut fungi parasitizing the *S. nutans* complex (Fig 5C). Within *Microbotryum* strains parasitizing *S. nutans*, we found no footprint of recombination between strains of the two main genetic clusters (Fig 5C) which was consistent with a scenario of divergence with ancient migration and no contemporary gene flow inferred from ABC demographic inferences.

## Discussion

We found that genetic structure and divergence history of anther-smut fungi parasitizing the *S. nutans* plant species complex closely mirrored those of their host plant genetic lineages. Such a congruence in population structure and divergence history of the host and the pathogen in the *S. nutans* complex, the distribution area of the genetic clusters in Western Europe, as well as the inferred dates of divergence suggest that the host and the pathogen differentiated in parallel following isolation in shared glacial refugia. To confirm the co-divergence events between *S. nutans* lineages and anther-smut fungi genetic clusters, we need to more accurately date the node of lineage divergence events in the host and the pathogen. Such molecular dating is essential to support co-divergence events as preferential shifts of the pathogen to closely related host species can also produce congruent phylogenies (de Vienne et al., 2013), but is currently difficult due lack of appropriate calibration points.

Nevertheless, our results rule out the hypothesis that anther-smut fungi on some of the *S. nutans* genetic lineages may result from specialization by host shifts from distant anther-smut fungi parasitizing other *Silene* species in sympatry or parapatry. This contrasts with the prevalence of host shifts between distant lineages observed in the Caryophyllaceae-*Microbotryum* system at larger evolutionary scales (Refrégier et al., 2008), but is consistent with the strong congruence reported between the population genetic structures of *S. latifolia* and its anther-smut fungi, also corresponding to glacial refugia footprints (Feurtey et al., 2016). Our study thus contributes to gain general insights into the processes of divergence in host-pathogen systems. While at large evolutionary scales, host shifts seem to be the rule (de Vienne, Hood, & Giraud, 2009; Thines, 2019), finer and more recent population subdivisions may more often result from codivergence due to shared geographic and climatic constraints, such as glacial refugia. The discrepancy between the two evolutionary scales may be due to recurrent pathogen lineage extinctions followed by recolonizations through host shifts: codivergence may occur frequently, but pathogen lineages would regularly go extinct over longer evolutionary scales and plant lineages would be recolonized by host shifts. This novel hypothesis can be tested by studying further closely related pairs of natural host-pathogen associations, which unfortunately still remain scarce.

The significant IBD pattern in both the *S. nutans* plant and anther-smut fungi and the correlation between the genetic distances between host and pathogen pairs when controlling for IBD suggest that the anther-smut fungi followed similar recolonization routes as the plant and/or became specialized on the host genetic lineages. However, contrary to the system *S. latifolia* - *M. lychnidis-dioicae* (Feurtey et al., 2016) and other host-pathogen systems (Barrett et al., 2008; Criscione, Poulin, & Blouin, 2005; Nieberding et al., 2008; Nieberding & Olivieri, 2007), we found higher levels of subdivision in *S. nutans* than in its associated anther-smut fungi. The weaker genetic structure in anther smut fungi compared to its host suggests again extinction of the pathogen lineage in a plant lineage, followed by recolonization from another fungal lineage, or the ability of an anther-smut lineage to remain generalist on two closely related plant lineages.

Future studies using cross-inoculation experiments between strains and host plants of different genetic clusters within the *S. nutans* complex could assess whether the congruence of host-pathogen genetic structure is associated with a pattern of local adaptation of the pathogen or of the pathogen, as in the system *S. latifolia* - *M. lychnidis-dioicae* (Delmotte et al., 1999; Feurtey et al., 2016; Kaltz et al., 1999). We could not obtain enough fresh material for this study to be able to test local adaptation. Furthermore, strong asymmetric postzygotic reproductive isolation was found between the Eastern and Western *S. nutans* plant lineages (Martin et al., 2017) and it would be interesting in future studies to explore if a similar reproductive isolation pattern is present between the Eastern and Western fungal lineages on *S. nutans*, and whether reproductive isolation can be found among anther-smut fungi.

We found clear differentiation between fungal populations parasitizing the two closely related *S. nutans* and *S. italica* species, supporting previous findings of differentiation based on a few strains and loci (Bucheli et al., 2000). No strong population structure was found within *S. italica* or within its associated anther-smut fungi. However, future studies with larger sampling distribution in the *S. italica* range in the Mediterranean Basin (Naciri, Pasquier, Lundberg, Jeanmonod, & Oxelman, 2017) may identify population differentiation. Cross-species disease transmissions, i.e. spill-overs of fungal strains, were more frequent on *S. nutans* than on *S. italica*, which may be due to unequal sampling size between the two *Silene* species or to biological differences. Some Caryophyllaceae plants indeed seem more susceptible to cross-species transmissions (Antonovics et al., 2002; de Vienne, Hood, et al., 2009; Hood et al., 2019). Putative spill-overs may actually correspond to other *Microbotryum* species, that were too rare to be identified in our population analyses using microsatellite markers, as revealed with the whole genome sequence data. In fact, two strains collected on *S. nutans* were actually assigned, based on whole genome sequences, to *M. lychnidis-dioicae,* specialized on *S. latifolia*.

The statistical comparison of demographic models and the whole-genome SNP analyses indicated the absence of recent gene flow between *Microbotryum* species parasitizing closely related *Silene* species, as well as between the Eastern and Western fungal lineages parasitizing *S. nutans*, although some spill-over cases were detected. Noteworthy, we also found no evidence of hybrid individuals between the two host plants species *S. nutans* and *S. italica* based on SNP data. The admixed barplots observed in the microsatellite STRUCTURE analyses must therefore have been due to low power of assignment rather than recent hybridization. This result was consistent with previous studies on other several anther-smut fungi, in which no hybrids were detected in natural populations, even between very closely related species that can hybridize in the laboratory (Abbate et al., 2018; Badouin et al., 2017; Gladieux et al., 2013; Petit et al., 2017). The only case where hybrids were detected was among closely related anther-smut fungi with overlapping host and geographic ranges, on *Dianthus* species (Petit et al., 2017). Ecological factors, such as different habitats, pollinator guilds or flowering time of the host species (Jürgens, Witt, & Gottsberger, 1996; Kephart, Reynolds, Rutter, Fenster, & Dudash, 2006), may constitute pre-zygotic factors favoring reproductive isolation of host-specialized anther-smut fungi in natural populations. Both extrinsinc and intrinsic post-zygotic barriers can be strong in anther-smut fungi (Giraud & Gourbière, 2012; Le Gac, Hood, & Giraud, 2007). Strong host specialization may indeed play a role in reproductive isolation, through migrant inviability and hybrid maladaptation on parental hosts, especially given the life cycle of *Microbotryum* fungi, with many spores falling on a plant and competing for systemic infection, and selfing being frequent, exposing hybrids to systematic competition with non-hybrids (Gibson, Hood, & Giraud, 2012). In addition, comparative genomics of anther-smut fungi showed presence of large genomic rearrangements and gene content variation between species (Branco et al., 2018; Hartmann et al., 2018), and experimental crosses suggested high frequency of hybrid sterility and abnormal genomic contents in hybrids (de Vienne, Refrégier, et al., 2009).

The low levels of gene flow among anther-smut fungi parasitizing different hosts found in *Microbotryum* fungi stand in high contrast with frequent reports of signatures of introgression in other fungal pathogens, such as crop pathogens or human disease-associated pathogens (Feurtey & Stukenbrock, 2018). Few studies have focused on fungal pathogens diversification in natural host communities while several evolutionary processes, such as time scale of divergence, host density and heterogeneity, are likely very different than those occurring on human-modified environment pathogens (Laine, 2005; Laine, Barrès, Numminen, & Siren, 2019; Stukenbrock & McDonald, 2008). To understand how biodiversity arises and what the mechanisms of host-pathogen evolution are over large evolutionary scales, we need more studies on co-evolutionary histories of parasites and their hosts in natural ecosystems. Studies of population genetic structure and divergence in plant fungal pathogens indeed remain so far mostly focused on crop pathogens (Barrès et al., 2008; Enjalbert, Duan, Leconte, HovmøLler, & De Vallavieille-Pope, 2005; Fournier & Giraud, 2008; Linde, Zhan, & McDonald, 2002; Saleh, Milazzo, Adreit, Fournier, & Tharreau, 2014; Stukenbrock, Banke, & McDonald, 2006; Zaffarano, McDonald, & Linde, 2008), in which patterns are heavily impacted by host genetic homogeneity and high abundance, as well as by human-mediated plant and pathogen dispersal. We found here strong population structure congruence between hosts and pathogens, in agreement with a previous study on other anther-smut lineages (Feurtey et al., 2016). Further studies on other natural systems are needed to assess whether this represents a general pattern in natural pathogen-host communities.

## Acknowledgements

We thank all the collectors cited in Table S1 and S7 for their contribution to plant material sampling, the Département de la Nature et des Forêts (Service Public de Wallonie, Belgium), B. Clesse (Centre Marie-Victorin, Cercles des Naturalistes de Belgique asbl), Ardenne & Gaume and Natagora for access to natural sites and for the authorization to collect plant material. We are grateful to the INRA MIGALE bioinformatics platform (http://migale.jouy.inra.fr) for providing computational resources and the Genotoul platform for sequencing. We thank the Plateforme de Genotypage GENTYANE INRA UMR1095 for help with microsatellite genotyping. This work was supported by the European Research Council (ERC) (starting grant GenomeFun 309403), the French National Research Agency (ANR) (Gandalf ANR Grant ANR-12-ADAP-0009), the Louis D. Foundation (Institut de France) to TG, and a Marie Curie European grant (PRESTIGE-2016-4-0013) to FEH. PT and CG thank the Région Hauts-de-France, and the Ministère de l’Enseignement Supérieur et de la Recherche (CPER Climibio), and the European Fund for Regional Economic Development for their financial support.

## Datasets

dataset 1: Hartmann F.E., Snirc A., Cornille A., Godé C., Touzet P., Van Rossum F., Fournier E., Le Prieur S., Shykoff J., Giraud T. 2019. Plant microsatellite and chloroplastic genotypes. Dryad doi (to be completed upon acceptance).

dataset 2: Hartmann F.E., Snirc A., Cornille A., Godé C., Touzet P., Van Rossum F., Fournier E., Le Prieur S., Shykoff J., Giraud T. 2019. Fungal microsatellite genotypes. Dryad doi (to be completed upon acceptance).

dataset 3: Hartmann F.E., Snirc A., Cornille A., Godé C., Touzet P., Van Rossum F., Fournier E., Le Prieur S., Shykoff J., Giraud T. 2019. Whole genome raw data of *Microbotryum* fungi infecting *S. nutans* and *S. italica*. NCBI Sequence Read Archive (SRA). BioProject xxx (ID available upon acceptance).

## Data accessibility

The plant and fungal microsatellite and chloroplastic genotypes are available at dryad XXX (to be completed upon acceptance). We deposited the genome raw data at NCBI Sequence Read Archive (SRA) under the BioProject xxx (ID available upon acceptance).

## Author contributions

TG and FH conceived and designed the project with the help of FVR. TG, PT, FVR, JS and EF collected samples. AS, CG, PT, SLP and FH genotyped the samples. FH and AC performed the analyses. FH, TG and FVR wrote the manuscript. All authors read and approved the final version of the manuscript and declare no conflict of interest.

## Supporting information

Additional Supporting Information may be found in the online version of this article.

